# Collective behavior emerges from genetically controlled simple behavioral motifs in zebrafish

**DOI:** 10.1101/2021.03.03.433803

**Authors:** Ariel C. Aspiras, Roy Harpaz, Sydney Chambule, Sierra Tseng, Florian Engert, Mark C. Fishman, Armin Bahl

## Abstract

Since Darwin, coordinated movement of animal groups has been believed to be essential to species survival, but it is not understood how changes in the genetic makeup of individuals might alter behavior of the collective. Here we find that even at the early larval stage, zebrafish regulate their proximity and alignment with each other. Two simple visual responses, one that measures relative visual field occupancy and the other global visual motion, suffice to account for the group behavior that emerges. We analyze how mutations in genes known to affect social behavior of humans perturb these simple reflexes in larval zebrafish and thereby affect their collective behaviors. We use model simulations to show that changes in reflexive responses of individual mutant animals predict well the distinctive collective patterns that emerge in a group. Hence group behaviors reflect in part genetically defined primitive sensorimotor “motifs”, which are evident even in young larvae.

## Introduction

Collective movements of animal groups are critical for species survival^1^. They provide protection from predation^2,3^, improve foraging^4,5^, and help with better energy utilization^6,7^. Presumably, the wide variety of group behaviors that characterize different species reflect underlying genetic changes, but to identify the precise role that individual genes play in controlling behavior, we need to first understand the underlying sensorimotor algorithms implemented by individual animals that give rise to emergent collective behavioral patterns.

Zebrafish, like many other aquatic animals, display a variety of distinct group behaviors, including shoaling, where individuals swim in proximity, and schooling, where all members of the group move in the same direction. To achieve such synchronized movements in groups, individual members need to assess certain properties of their near neighbors, including their speed, distance, and orientation, and they need to respond rapidly to these features and then execute the appropriate motor commands. The ability to perform such socially relevant sensorimotor transformations, and thereby the ability to form groups, differ between different genetic backgrounds^8–11^ and are modified by hunger^12–14^ and innate “personalities’’ of individual fish^15,16,17,18^. Further, while inputs from several sensory modalities, such as lateral line mechanoreception^19,20^, olfaction^21,22^ and vision, all likely play a role in this process, vision is critical to certain attributes, such as the rapidity of turning responses, the necessary integration of distal cues, and the precision of the alignment responses^11,23,24^. Mapping out the algorithmic rules and neurophysiology underlying collective behaviors can be more readily accomplished in larval zebrafish when the brain is transparent and circuits are simpler than in adults^25^. However, while reflexive responses to stimuli emanating from conspecifics have been described in various contexts^19,20^, there has been no evidence of either shoaling or schooling-like behavior in zebrafish larvae younger than ∼20 days post fertilization (dpf)^26,17^.

Here we identify two visual reflexes that are present from 7 dpf and are predictive of the emergent patterns of the collective as fish mature. First, young larvae repel away from regions of high visual clutter, leading to a dispersal of the group. At later developmental stages, this dispersal reflex shifts to attraction and aggregation behaviors. Second, larvae display a strong tendency to move along with whole field motion stimuli, a well-described behavior known as the optomotor reflex (OMR). When individuals swimming within a group use their neighbors’ swimming direction as motion cues, this reflex leads to an emergence of mutual alignment between close neighbors and induces collective motion of the whole group. The combined developmental maturation of both reflexes can then explain emergent shoaling and schooling behavior.

Mutations in genes associated with autism and schizophrenia quantitatively alter these two visuomotor responses, and these changes are predictive of the distinct emergent behaviors of groups of the mutant fish. Thus, subtle alterations in simple behavioral motifs of the individual can account for complex emergent patterns of groups.

## Results

### Visually driven aggregation and alignment in larval and juvenile zebrafish

We analyzed behavior of wild-type larval zebrafish in groups of five, swimming together in a small arena, where an overhead camera is used to monitor the position, orientation, and speed of each individual over extended time periods (**Fig. 1a**, methods).

**Figure 1.**
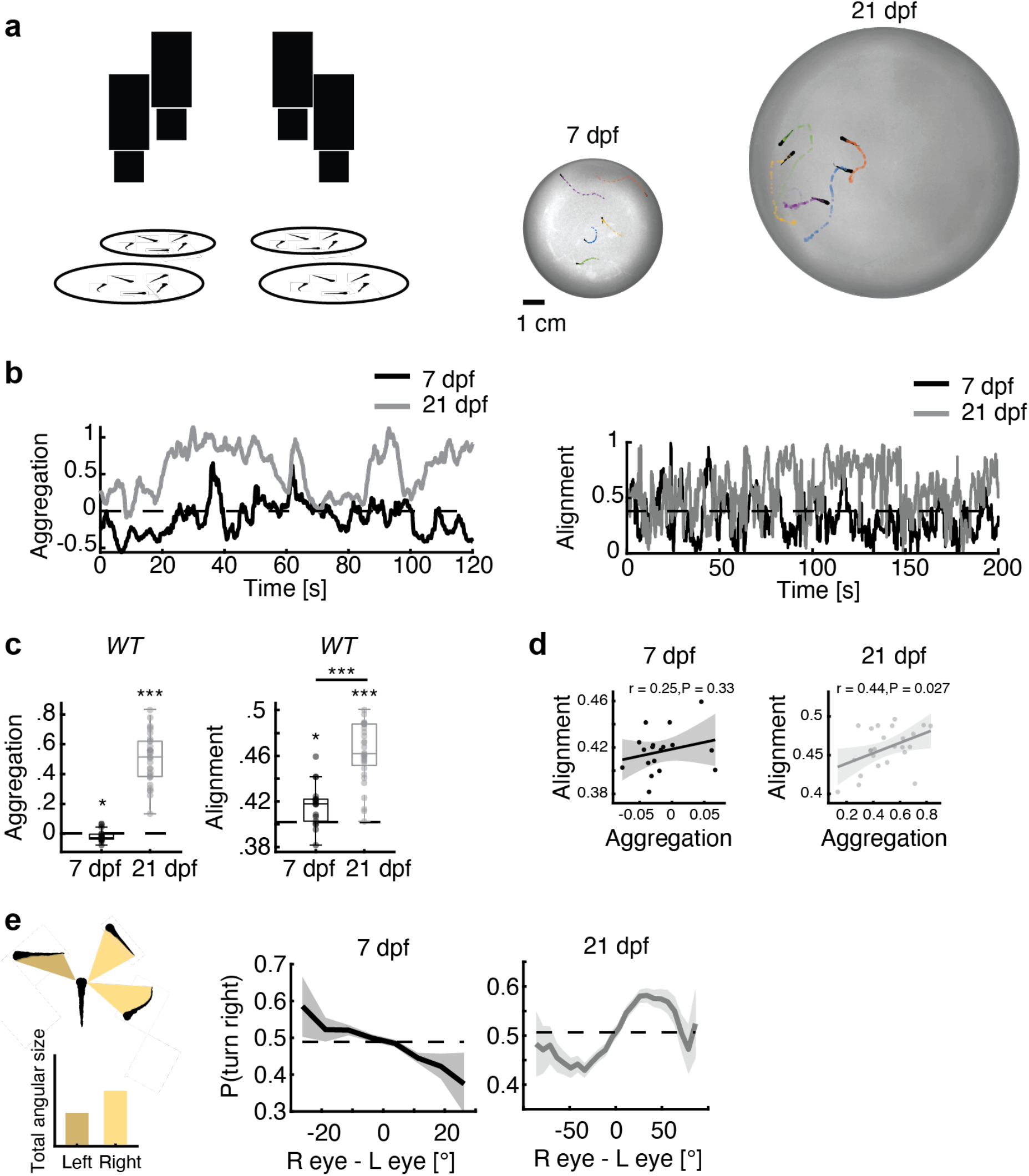
Aggregation in developing zebrafish. **(a)** Left: Groups of five fish were tested in circular arenas, while overhead cameras recorded their behaviors. Right: Position and body orientations of each fish were extracted from the movies. (**b)** Example traces of the aggregation index (left) and alignment index (right) (see methods) of groups of 7 and 21 dpf wild type fish. Dotted lines represent chance levels. 21 dpf fish show higher aggregation and alignment than 7 dpf fish. (**c**) Left: at 7 dpf fish are significantly less aggregated than expected by chance (P<0.05, N=16 groups; t-test), while 21 dpf fish form tight groups (P<0.001, N = 25 groups; t-test). Right: 7 dpf fish are more aligned than expected by chance (P<0.005, N = 18 groups; ot-test) and 21 dpf fish are more aligned than 7 dpf fish (P<0.001, t-test). **(d)** Correlation of alignment and aggregation in 7 (left) and 21 dpf (right) fish. Significant correlation emerges for older larvae. (**e**) Effect of “visual clutter”. Left: we reconstruct the visual angle that each neighboring fish is expected to cast on the retina of a focal fish (Methods). Right: The difference between total angular area (or “visual clutter”) experienced by each eye significantly modulates the probability to turn away (7 dpf) or towards (21 dpf) the more cluttered visual field

We find that as animals mature from 7 dpf to 21 dpf, they swim more closely to one another, are more aligned, and exhibit faster swimming speeds (**Fig. 1a-c, Fig. S1a**). In addition, at 21 dpf a positive correlation between mutual alignment and aggregation starts to emerge (**Fig. 1d**), a phenomenon that suggests conspecifics swimming closer to each other evoke stronger alignment responses than do more distant fish. Even at 7 dpf, groups already exhibit evidence of interactions. At this time, however, aggregation indices are distinctly smaller than chance, indicating that young larvae display mutual repulsion rather than attraction. Alignment is slightly enhanced when compared to chance. To explore the algorithmic basis of the transition from dispersed to aggregated groups, we analyzed the turning behavior of fish in response to the occupancy of their right and left visual fields, i.e. the retinal “clutter” generated by the presence of conspecifics swimming in the vicinity^27^ **(Fig. 1e**, left). We find that 7 dpf fish tend to turn away from the more highly cluttered area, while 21 dpf animals turn towards it (**Fig. 1e**). This simple visuomotor response to difference in retinal clutter will lead to a dispersal (7 dpf) or aggregation (21 dpf) phenotype, depending on the sign of the transformation.

We next tested whether perturbations through targeted genetic mutations will cause specific changes in the clutter avoidance curves (**Fig. 1e**), and thereby generate quantitative changes in aggregation indices and the associated shoaling phenotype in mutant animals. To that end, we selected fish with mutations in two genes that have been shown to generate specific social phenotypes in adults^11^. The first gene, *scn1lab*, codes for a sodium channel. Its mutation is associated with Dravet syndrome in humans, and causes more scattered group behaviors in adult zebrafish. We evaluated heterozygous fish of two alleles (*scn1laball*_*ele1*_ and *scn1lab*_*allele2*_) because homozygous mutations are early lethal. The second gene, *disrupted-in-schizophrenia (disc1)*, encodes a scaffolding protein associated with schizophrenia in humans, and zebrafish with homozygous deficiency in *disc1* display increased group cohesion as adults^11^. We find that larvae with mutations in both genes behave differently from wild-type fish in the group swimming assay. At 7 dpf, both *scn1lab* mutants show an increase in dispersal when compared to their wild-type siblings, which manifests as a reduced aggregation index (**Fig. 2a, S1b**). 21 dpf *scn1lab* mutants switch from dispersal to aggregation, as do their wild-type siblings, but their increase in aggregation is much more moderate (**Fig. 2a, S1b**), which is in line with an enhanced mutant dispersion phenotype also at later developmental stages. In contrast, 7 dpf *disc1*^-/-^ fish show a small but significant increase in aggregation relative to their *disc1*^*+/+*^ sibling controls, an effect that becomes much more pronounced by 21 dpf (**Fig. 2b**).

**Figure 2.**
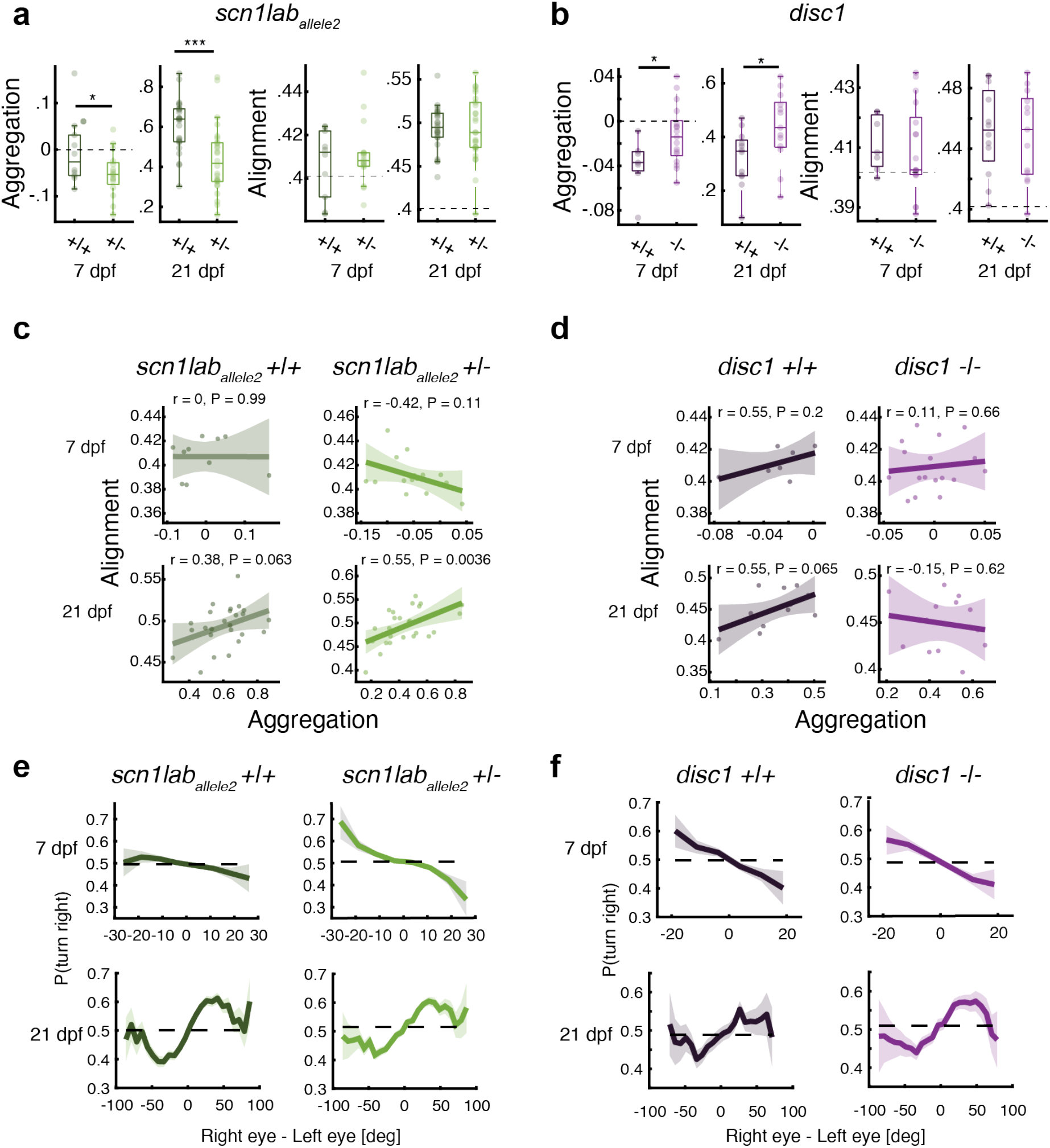
Single gene mutations affect aggregation and alignment of developing zebrafish. In all panels, +/+ refers to fish that are wild-type at the mutant locus, and that are siblings of the mutant fish. **(a)** Left: at 7 dpf, mutant *scn1lab* fish are significantly less aggregated than wild-type siblings (P<0.05, N=26 groups; ttest) and are also less aggregated than expected by chance (or, more dispersed) (P<0.001, N=16 groups; ttest). Dotted lines represent chance levels. At 21 dpf, fish are more aggregated, yet *scn1lab*^*+/-*^ fish aggregate less than *scn1lab*^*+/+*^ fish (P<0.001, N = 51 groups; ttest). Right: 7 dpf *scn1lab*^*+/-*^ are more aligned than expected by chance (P<0.05, N = 16 groups; ttest) and 21 dpf fish are more highly aligned than 7 dpf fish. No direct effects of the mutations were observed at either age. (**b**) Left: at 7 dpf, mutant *disc1* fish are more aggregated (less dispersed) than sibling controls (P<0.05, N=22 groups; ttest). At 21 dpf, fish are more aggregated, and here again mutant *disc1* fish show an increase in aggregation compared to sibling controls (P<0.05, N = 23 groups; ttest). Right: 21 dpf fish are more aligned than 7 dpf fish, but no effect was observed for the *disc1* mutation. **(c)** Correlation of alignment and aggregation in *scn1lab*_*allele2*_ 7 (top row) and 21 dpf (bottom row) fish. No correlation is observed at 7 dpf (Top), while in 21 dpf *scn1lab*_*allele2*_^+/-^ fish (Bottom right) show the strongest correlation of aggregation with alignment. **(d)** No correlation of alignment and aggregation is observed in either *disc1*^*-/-*^ or *disc1*^*+/+*^ *at* 7 dpf, while wild-type sibling controls show a marginally significant correlation at 21 dpf. **(e)** *scn1lab*_*allele2*_^+/-^ fish show a significant increase in the tendency to turn away from visual clutter at 7 dpf, whereas at 21 dpf *scn1lab*_*allele2*_^+/-^ fish show a slightly reduced tendency to turn towards clutter. (**f)** mutant *disc1-/-* fish show no clear difference compared to wild-type siblings at either age.

When we specifically compared the clutter response curves between *scn1lab*^+/-^ fish and wild-type siblings, we found a significantly enhanced tendency to turn away from high clutter in 7 dpf mutants, and a slightly reduced tendency to turn towards high clutter in 21 dpf fish (**Fig. 2e**). This is in agreement with the observation that both ages display smaller aggregation indices when compared to their wild-type siblings. The *disc1*^-/-^ mutation, on the other hand, displays a slight “flattening” at the edges of the clutter response curve at 7 dpf, and a similarly small enhancement of the tendency to turn towards high visual clutter at 21 dpf (**Fig. 2f**). These trends are both qualitatively predictive of the increase in aggregation indices at both ages (**Fig. 2b**). However, uncovering the precise relationship between the tuning curves and aggregation indices requires a more detailed and quantitative analysis, and our approach using generative models towards this end is described below (**Fig. 4**).

Both strains of mutant fish, as well as their wild-type siblings, show significantly enhanced alignment by 21 dpf compared to 7 dpf, and no significant difference in alignment was apparent using this assay between mutant animals and wild-types at either age (**Fig. 2a**,**b**). We do, however, find that the correlation of alignment with aggregation is strongest in *scn1lab*^*+/-*^ 21 dpf mutants (in both alleles) (**Fig. 2c, S1d**), and it is absent in 21 dpf *disc1*^*-/-*^ animals (**Fig. 2d**). This observation suggests that mutations might cause subtle changes in alignment, perhaps dependent upon proximity, which we explore further using a more targeted approach to extract alignment phenotypes, as described below in the OMR experiments.

In summary, we find that conspecific fish do interact even at 7 dpf, as measured by a tendency to move away from visual clutter and by a higher-than-chance alignment index. This interaction precedes the well-described tendency to move towards clutter by 21 dpf. Notably, mutations associated with human social disorders have a detectable effect on behavior as early as 7 dpf, and the effects become more pronounced as animals mature.

### Mutant larval zebrafish show specific changes in their ability to align with motion

In addition to movement decisions that result in changes in proximity, the other key element of group behavior in fish is alignment, which gives rise to schooling behavior. We show that larval zebrafish, even as young as 7 dpf, already display a slight tendency to align with each other (**Fig. 1c**). The tendency to orient to the motion stimuli presented by other animals swimming in their vicinity can be quantitatively analysed using coherent-dot-based optomotor response (cdOMR) assays. In these assays animals reorient and align their swimming direction with a whole field visual motion stimulus presented in close proximity directly underneath the animal. Specifically, free swimming individuals, age 7 dpf, were presented with clouds of flickering small dots that drifted slowly either to the right or left relative to their body orientation (**Fig. 3a** and **Video S1**). We varied the degree of coherence, i.e. the fraction of dots moving in the same direction. This makes it more challenging to identify motion direction such that fish have to temporally integrate information to make appropriate swimming decisions^28,29^. These decisions were then quantified as a function of the coherence level by the fraction of swims following the direction of the motion stimulus (“probability correct”) and the time of quiescence between consecutive swims (“interbout interval”) (**Fig. 3b**,**c**). We also assessed the probability of swimming in the same direction for consecutive bouts, even when not stimulated by motion drift, to assess the tendency of larval zebrafish to repeat the same motor action over extended periods of time^30^ (**Fig. 3d**). We further quantified how swimming decisions for different coherence levels improve as a function of time (**Fig. S2b**) and characterized turn angle distributions (**Fig. S2c**).

**Figure 3.**
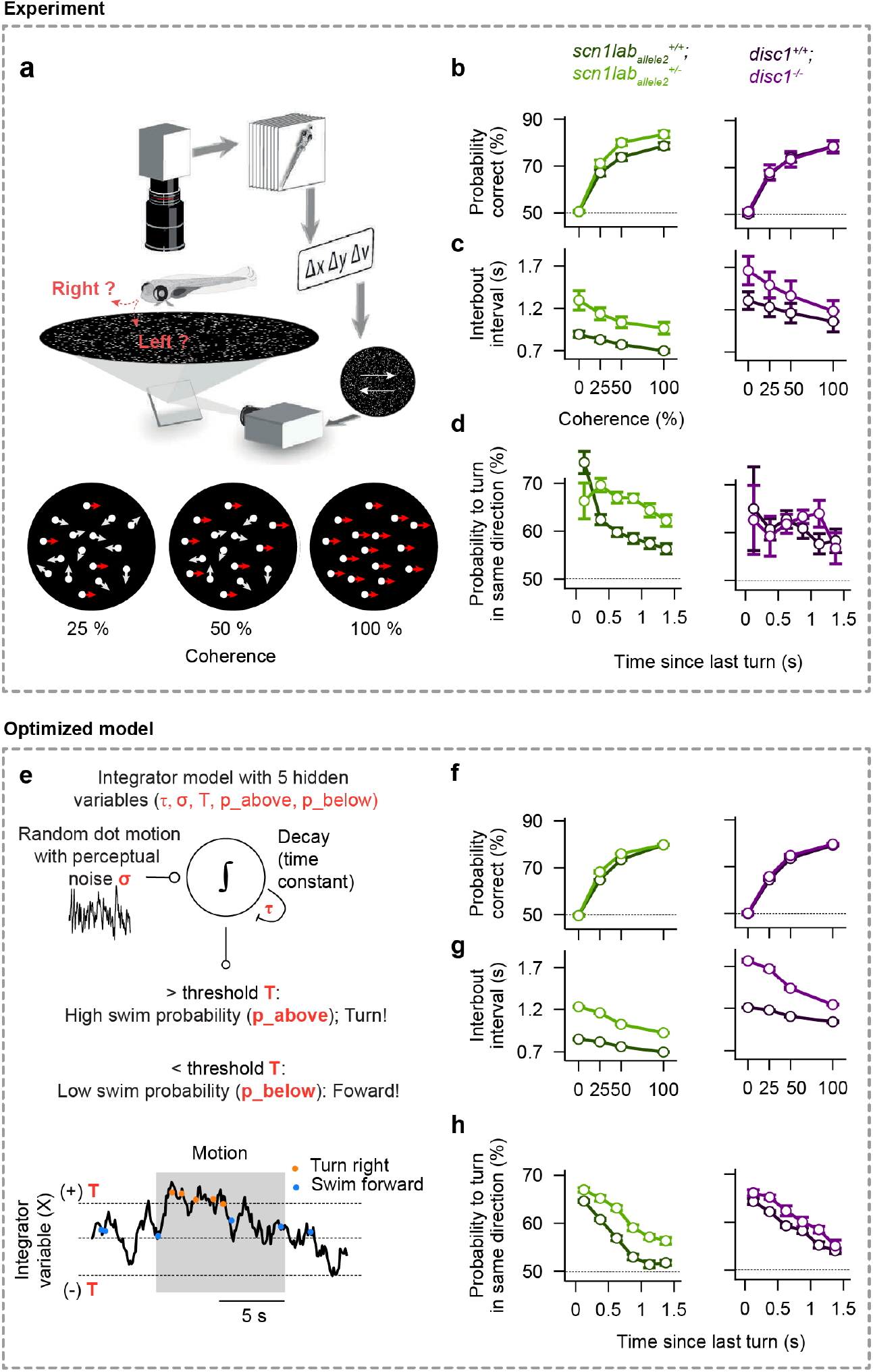
Mutant 7 dpf larval zebrafish display differential integration and alignment phenotypes, which can be quantitatively captured by a simple integrator model. **(a)** Experimental setup (adapted from ref. ^28^). A single larval zebrafish swims freely on top of a projected cloud of randomly moving dots. Dots move continuously at different coherence levels either to the right or left relative to the body orientation of the animal (**Video S1**). (**b**) The probability to correctly align with the coherent motion stimulus as a function of coherence strength. *Scn1lab*_*allele2*_^*+/-*^ mutant fish (bright green) show a performance improvement relative to *scn1lab*_*allele2*_^*+/+*^ wild-type sibling controls (dark green). The *disc1* mutant (magenta) shows no phenotype compared to respective sibling controls (dark magenta). (**c**) The interbout interval as a function of coherence. Values are elevated for both mutants relative to wild-type sibling controls. (**d**) The tendency to turn in the same direction as a function of the time since the last bout during randomly flickering 0% coherence stimulation. Responses are elevated for the *scn1lab*_*allele2*_ mutant but not for the *disc1* mutant, relative to wild-type sibling controls. (**e**) Integrator model with decision threshold (T), perceptual noise (σ), leak time constant (τ), and probabilities to make a turn or swim forward (p_above_, p_below_, depending on whether the integrated value is above or below the threshold). (**f–h**) Optimized model results, analyzed and displayed as in **b–d**. The model accurately captures the behavioral features of both wild-type and mutant larvae. N = 44, 36, 21, and 16 individually tested fish for genotypes *scn1lab*_*allele2*_ ^*+/+*^, *scn1lab*_*allele2*_ ^*+/-*^, *disc1*^*+/+*^, *and disc1*^*-/-*^, respectively, in **b–d**. N = 12 models (different optimization repeats) for each genotype in (**f–h**). Error bars in **b–d** and **f–h** are ±sem.

When we tested our mutant strains in this assay, we find that, compared to wild-type siblings, fish of both *scn1lab*^*+/-*^ alleles have an increased probability of responding correctly as a function of coherence, and that they respond with longer delays (**Fig. 3b** and **Fig. S2b**). They further display increased interbout intervals (**Fig. 3c**) and show an increased probability of turning in the same direction for consecutive bouts (**Fig. 3d**). The *disc1*^-/-^ mutants, on the other hand, differ only in having longer interbout intervals than their wild-type siblings (**Fig. 3c**). Turning distributions between mutant animals and sibling controls indicated a slight increase in turn angle, compared to controls, for all tested genotypes (**Fig. S2c**). Hence, mutations in both genes cause subtle differences in the animals’ ability to integrate information over time and to align with motion drift in their environment.

### Drift-Diffusion model for motion integration to explain alignment

We have previously shown that the responses of larval zebrafish to coherent dot motion can be well described by the computational framework of a “Drift Diffusion Model” (DDM)^31,32^ (**Fig. 3e**). This model uses only five parameters, the internal noise (**σ**), integration and decay time constants (**τ**), decision thresholds (**T**), and two swimming probabilities (**p_below** and **p_above**; see methods) to fully describe the behavior. Notably, none of these parameters can be measured directly through experiments. We therefore resorted to a multi-objective fitting approach which allowed us to automatically extract those values and systematically explore their variation due to the mutations (**Fig. S3, Fig. S4**, and methods). We show that this strategy quantitatively captures subtle behavioral features across the different mutants. For example, we find that, when compared to the respective sibling control animals, the threshold variable (T) decreases in *scn1lab* mutant fish, while it increases for the *disc1* mutant (**Fig. S4b**). Knowledge about these behavioral variables allows us to make specific predictions about corresponding neural circuit changes in mutant animals^28,29^ and further, these variables can provide the critical substrate model simulations of fish in more complex scenarios (see below). Thus, modeling wild-type and mutant fish behavior exclusively based on the DDM with these extracted variables allowed us to test whether this framework is sufficient to explain the behavioral results. We find that the model predicts most of the experimental data (**Fig. 3b–d,f–h, Fig. S3b**,**c**,**e**,**f**). This suggests that the DDM provides an adequate framework to quantitatively describe alignment behavior in groups, and that it is capable of reliably extracting hidden integration and decision-making variables in mutant animals.

### Models based on two simple reflexes explain emergent collective behavior

We tested whether the two basic reflexes, i.e. the “clutter response” (**Fig. 1e**) and the coherent moving dot response (cdOMR) (**Fig. 3a**), are sufficient when applied to individual fish, to explain the emergent dynamics of zebrafish shoaling and schooling behaviors. To that end, we simulated groups of five virtual fish swimming in a circular arena, where each individual agent follows only the computations predicted by our two assays (**Fig. 4a**). The “clutter response computation” is the measurement of the clutter projected by the four conspecifics onto the left and right eye. The “coherent moving dot computation” measures the perpendicular motion component generated by all other fish. Both signals are integrated over time and compared to the threshold, allowing the model to make decisions about whether to move forward or make turns, as described by the DDM model (**Fig. 3e** and methods).

**Figure 4.**
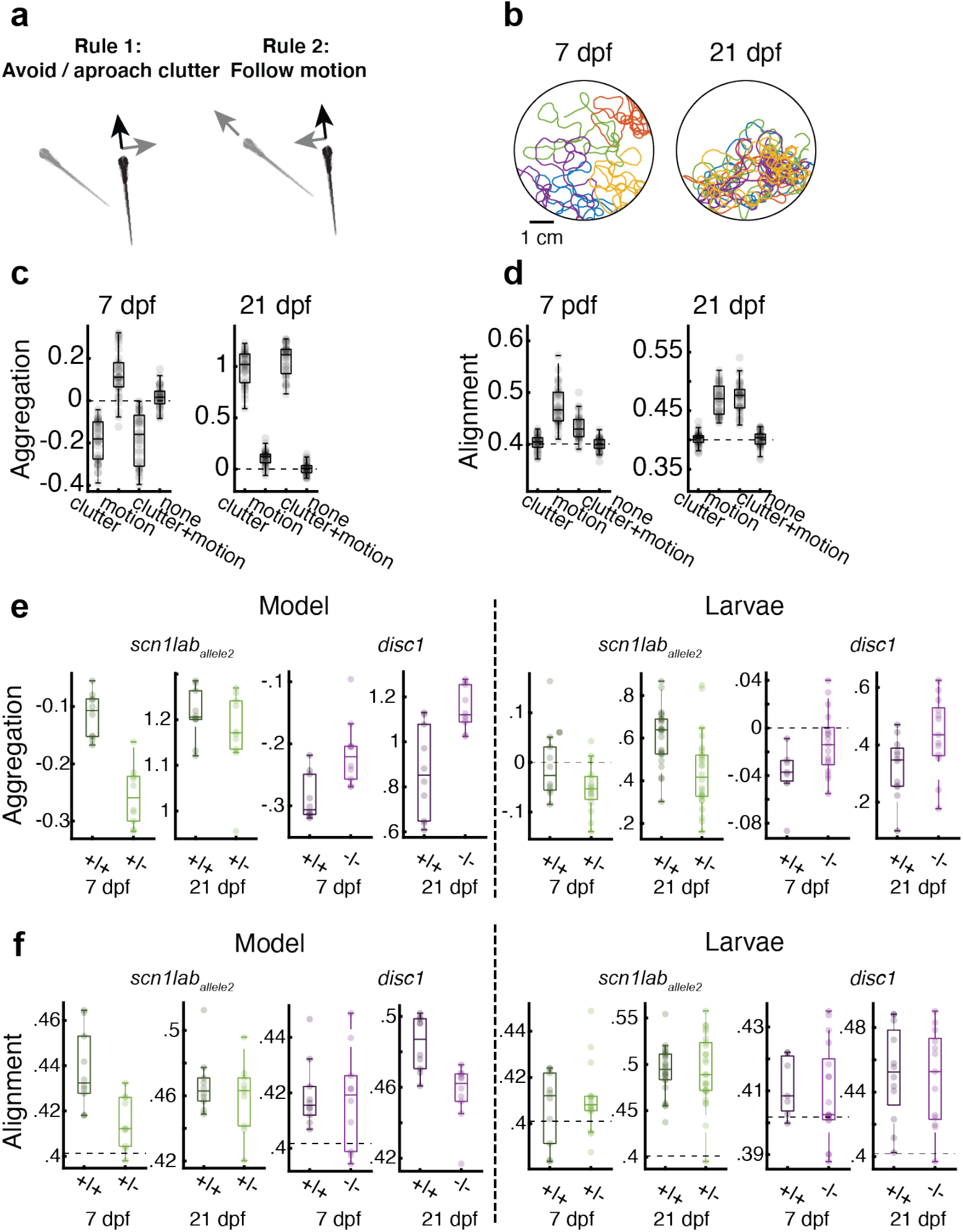
Simple visuomotor reflexes qualitatively predict emergent group behavior across genotypes and development. **(a)** Schematic of the model, using only two simple algorithmic rules: First, fish are repelled or attracted to visual clutter (see **Fig. 1e**). Second, fish use global motion cues for turning decisions (see **Fig. 2a**,**e**). Both cues are integrated over time and evaluated according to our integrator model (**Fig. 2e**). Model parameters for the clutter response strength are directly extracted from group swimming experiments (**Fig. 2e,f**). Model parameters for the integration and decision-making process are taken from our multi-objective parameter fitting results (**Fig. S4b**). The model is hence almost fully constrained by experimental data. (**b**) Example trajectories of simulated wild-type 7 dpf and 21 dpf, based on the rules shown in **a**, showing that older fish start to aggregate and align with one another. (**c**,**d**) Aggregation and alignment for wild-type simulations for all possible rule combinations (neither rule, only motion, only clutter, or both rules) for 7 dpf and 21 dpf model fish. Alignment does not emerge with attraction alone, it additionally requires animals to perform motion integration. Conversely, motion integration induces some level of aggregation. Parameters for wild-type animals are the same as for the sibling controls for all tested mutant lines. (**e**,**f**) Aggregation and alignment for 7 dpf and 21 dpf mutant model fish with respective sibling controls (left) and the corresponding data of real fish (right, same data as in **Fig. 2)**. Our model can qualitatively predict all group behavior phenotypes across ages as found in our experimental group assay (**Fig. 2a,b**). N = 36 model simulations (each model uses different parameter sets, following our repeated model optimization, **Fig. S4b**) in (**c**,**d**) and N = 12 model simulations for each genotype in (**e**,**f**). Experimental data in (**e**,**f**) is the same as in **Fig. 2a,b**).

Using these two computations, we simulated swim trajectories of groups of 7 dpf and 21 dpf “virtual” fish (**Fig. 4b, Videos S2** and **S3**), and extracted aggregation and alignment indices, as done for groups of real fish (**Fig. 1**). Running the simulations for the different experimentally obtained variables and response curves revealed that the model produces results that qualitatively match the experimental findings: 7 dpf wild-type virtual larvae show a tendency to repel each other (with slightly negative aggregation indices), while 21 dpf virtual animals show strong aggregation behavior (**Fig. 4c**). Alignment indices significantly increase from 7 dpf to 21 dpf, as observed in the experimental results (**Fig. 4d**). Since we used the same cdOMR variables for 7 dpf and 21 dpf simulations, this improvement in alignment is likely a consequence of the enhanced aggregation values in older animals, which leads to a more pronounced effect of visual motion cues and consequently stronger alignment **(Fig. 1d)**. To further, and explicitly, probe the interdependence of aggregation and alignment, we asked how well each rule by itself will predict aggregation and alignment indices respectively (**Fig. 4c**), and how they interact when combined. We find that aggregation indices are, as expected, dominated by the clutter response rule (although the addition of the motion response rule slightly enhances the tendency to aggregate). Alignment indices, on the other hand, which depend predominantly on the motion response rule, are significantly modulated and brought into far better agreement with observed data when the clutter response is added. Thus, the combination of the two attributes, clutter response and dot motion response, predicts that the reduced aggregation of 7 dpf fish will lead to weaker alignment, and the strong aggregation in 21 dpf animals will lead to better alignment, both results that are also apparent in the experimental data.

We next ran the complete simulation, combining both clutter and motion computations, using the model parameters that we extracted from our experimental assays for each genotype (**Fig. 2e-f, Fig. S1e**, and **Fig. S4b**). Here we find, also in agreement with the experimental data, that the tendency of 7 dpf wild-type virtual larvae to repel each other is enhanced in *scn1lab*^+/-^ mutant fish, and diminished in *disc1*^-/-^ animals (**Fig. 4e**). This phenotype carries robustly into 21 dpf animals, where *scn1lab*^+/-^ show a decreased and *disc1*^-/-^ an increased aggregation phenotype compared to wild-type virtual siblings. Moreover, our model reproduces that both 7 dpf and 21 dpf mutant animals show minimal alignment phenotypes compared to sibling controls (**Fig. 4f**).

The success of this straightforward minimal model in reproducing the experimental results suggests that, at least in the larval animal, genetic effects upon just the two visual responses suffice to predict most attributes of the emergent behavior of the group.

## Discussion

Here we find that even very young larval zebrafish interact with each other, and that their group dynamics are well predicted by two visual responses: the retinal clutter response and the OMR. These two visuomotor assays explore the tendency of fish to attract and to align with each other, key attributes of collective behaviors known as shoaling and schooling. We show that mutations in genes associated with autism and schizophrenia alter these two visual responses in subtle ways and that these changes are qualitatively predictive of emergent mutant shoaling and schooling phenotypes. Importantly, these subtle effects are already detected at fish as young as 7 dpf, and carry over to the adult fish.

Specifically, mutual attraction is predicted by the tendency of larval fish to turn and reorient based on the relative retinal occupancy, or retinal clutter, between the two eyes. Alignment, on the other hand, is predominantly predicted by the OMR, where larval fish reorient and turn to swim along with whole field motion patterns in their surrounding. In theory, with appropriate parameter choice, alignment can also emerge purely based on clutter-based algorithms^24^. However, given our experimental and modeling results, we believe that for larval zebrafish the clutter response is not sufficient to induce alignment but that it can serve to enhance it when combined with whole field motion stimuli.

We use here a modified OMR stimulus of coherently moving dots with short lifetimes, which adds components of temporal and spatial integration to the assay, and permits modeling of latent, or hidden, variables in wild-type and mutant fish. Because 7 dpf larvae do not aggregate into shoals or seek the vicinity of other fish^17,26,33^, any responses to conspecific stimulation that in past studies have been observed at this early age^20,19^, were assumed to be unrelated to shoaling or schooling behavior. Here we find that larval zebrafish repel, rather than attract, each other at this young age, which leads to a distinct global dispersion - or negative aggregation - phenotype. This dispersal can be explained by a clutter response of negative sign, where fish move away from the side with greater clutter rather than towards it. The effect is modest so it might have been missed for these reasons at young ages. We find that this repulsion phenotype switches to attraction with age, so that by 21 dpf the animals tend to form more familiar aggregates and shoals^33,26^. Because 7 dpf larvae are not very motile and tend to live in protected areas with little water flow, this repulsion might assure them of sufficient oxygenation from relatively unstirred surroundings^34^, and it might help them avoid frequent collisions in the cramped quarters typical for densely populated clutches. The switch to aggregation at 21 dpf likely helps keep groups of older animals together when they start exploring larger areas of their environment and when they begin migrating over longer distances.

Alignment is critical for adult schooling behavior. We show here that alignment in free-swimming groups of fish, starts to emerge around 7 dpf, and becomes more robust by 21 dpf. While no significant differences in alignment indices are apparent between any genotypes in the free-swimming groups, we find that 21 dpf scn1lab^+/-^ mutant fish show the strongest correlation of alignment with proximity in all strains tested (**Fig. 2c**), suggesting that these mutants might be more responsive to nearby moving stimuli. Indeed, when tested with the coherent dot assay, we find that *scn1lab*^*+/-*^ mutant fish align more accurately with moving dots than do their wild-type siblings, suggesting that the coherent-dot OMR can serve as a powerful tool to quantitatively dissect and uncover subtle alignment phenotypes that are difficult to extract in group swimming assays. This is concordant with a separate study in adult zebrafish, where it was found that *scn1lab*^*+/-*^ adults also exhibit enhanced alignment when swimming in close proximity^11^.

Our model, which is based solely on these two simple visuomotor transformations, can account for a large fraction of the complex collective interactions that occur in groups. The model’s simplicity and accuracy, and its applicability even to 7 dpf larvae with their relatively simple and accessible brains, makes it a practical entry-point to dissect the cellular nature of the algorithms that drive collective behaviors. Although fish have many sensory inputs that contribute to group behaviors, zebrafish are highly visual^28,30,35^, suggesting that visual drives likely play a dominant role. Other sensory modalities, such as somatosensation through the lateral line^19,20^ and olfaction^13,21^, undoubtedly also can play a role in modulating social interactions, as might currently less decipherable elements such as “internal state”^12– 14^ and “personality”^15–18^.

The specific genetic perturbations we have studied are in genes related to human psychiatric disorders. The human SCN1A gene (the orthologue of the zebrafish *scn1lab* gene) is associated with Dravet’s syndrome (where patients have epilepsy and developmental disorders including autism) and DISC1 is associated with schizophrenia. Interestingly, atypical visual reflexes, including the optokinetic response, have been noted in both autism and schizophrenia^36^, and there is evidence for linkage between such perceptual differences and social abnormalities in autism^37^. Perhaps reduction of complex behaviors to simple underlying reflexive motifs may help to characterize complex disorders. Quantitative characterisation of changes in these reflexes in mutant zebrafish facilitates analysis of the underlying cellular defects and enables screens for reparative therapeutics^38^.

## Acknowledgements

We thank Jacob D. Davidson, and Mark Fishman’s and Florian Engert’s lab members for discussion and advice. We thank Cristina Santoriello for technical and logistical support in addition to discussion and advice. We thank Kristian Herrera, Martin Haesemeyer and Katrin Vogt for critical reading of the manuscript and constructive feedback. This work was supported by a grant from Fidelity Biosciences Research Initiative and sponsored research support from NIBR to MCF. F.E. received funding from the National Institutes of Health (U19NS104653, R43 OD024879, 2R44OD024879), the National Science Foundation (IIS-1912293) and the Simons Foundation (SCGB 542973). A.B. was supported through the German Emmy Noether Program (BA 5923/1-1) and by the Zukunftskolleg Konstanz.

## Methods

### Zebrafish

To generate larvae for sibling controlled experiments, heterozygous fish were incrossed. For *scn1lab* experiments, the *scn1lab* ^*+/-*^ fish were crossed with AB wild-type. Clutches were raised in small groups (20-30) in 15 cm petri dishes with fish facility water in 14h light, 10 hr dark cycle at constant 28 °C. At 4 days post fertilization (dpf), larvae were fed rotifers or paramecia daily with 50% water change. Behavior experiments were done at 7 and 21 dpf. All experiments follow protocols approved by Harvard IACUC.

### Group assay

We used custom-designed experimental arenas of different sizes D=6.5,12.6 cm (for groups of 7 and 21 dpf fish), H = 1 cm made of 1/16” PETG plastic. Arenas had a flat bottom and curved walls to encourage fish to swim away from the walls and were sandblasted to prevent reflections. Every experimental arena was filmed using an overhead camera and lit from below using infrared light (same as in the dot motion assay). Images were acquired at ∼39 fps and were segmented online to separate fish images from the background. The segmented images were then analyzed offline to extract continuous tracks of the fish (^39^ for details of segmentation and tracking algorithms). All acquisition and online segmentation were performed using custom-designed software written in Matlab.

### Individual and group properties of free-swimming fish

We used the extracted center of mass position of every fish (*fish i*; 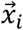) to calculate the velocity of the fish 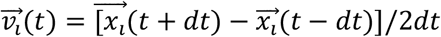, where dt is 1 frame or 0.025 s. The speed of the fish is then 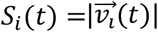, and the direction of motion is 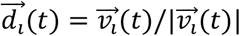.

For the group, we calculate a normalized measure of group aggregation: *Aggregation* =−*log*(*NN*_1_/*NN*_1_^*shuffled*^) where *NN*_1_ is the average nearest neighbor distance.*NN*_1_^*shuffled*^ is the same distance calculated from control groups created by shuffling fish between groups such that all fish in a shuffled group were chosen from different real groups. Positive aggregation values mean that real groups are more aggregated than shuffled controls and 0 means aggregation is at chance level. Group alignment was defined as 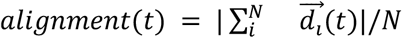, where N is the number of fish in the group and alignment value is bounded between 0 - all fish are pointing in different directions and 1 - all fish swim in the same direction.

### Estimating visual occupancy using ray casting

To estimate the visual angle that each neighbor in the group cast on the eye of a focal fish (*fish i*), we casted 1000 rays from each eye spanning 165°from the direction of motion towards the back of the fish, leaving a total of 30°of blind angle behind the fish. This amounts to an angular resolution of ∼0.165°per line. We then detected all pixel values representing fish in the paths of the rays and calculated the visual angle occupied by each fish and the total visual angle experienced by each eye (**Fig. 1e**).

### Motion assay

The assay has been described previously^28^. In brief, 7 dpf larvae were placed in custom-designed acrylic dishes (12 cm diameter, 5 cm height, black rim, and transparent bottom). The scene is illuminated from below with infrared LED panels (940 nm, Cop Security). The fish are tracked with a camera (Grasshopper 3), a zoom lens (Zoom 7000, 18-108 mm, Navitar), and a long-pass filter (R72, Hoya). Posture analysis is performed in real-time using custom-written software using Python 3.7 and OpenCV 4.1. Stimuli were presented from below (Aaxa P300 Pico Projector) onto mildly scattering parchment paper and consisted of ∼1000 small (2 mm) white dots on a black background. We showed 0% coherence as a baseline stimulus for 5 s and then switched to (25%, 50%, or 100%) coherent motion at a constant speed (1.8 cm/sec) for 10 s. Motion either went rightward or leftward relative to the body orientation of the fish. Each dot persisted for only 200 ms on average and stochastically disappeared and reappeared at a new location, so as to prevent fish from tracking individual dots. Following the coherent stimulus, the stimulus reverted to 0 % coherence baseline.

### Genotyping

For group assays (**Fig. 1, Fig. 2**, and **Fig. S1**), fish were genotyped at 2-3 dpf using Zebrafish Embryonic Genotyper (wFluidx) or fin clipping and high-resolution melt analysis (HRM; primer sequences in supplementary table1). Following all experiments, genotypes are confirmed by HRM following DNA extraction using hot shot genomic DNA preparation. Briefly, whole larvae are dissolved in 25 ul alkaline solution (25 nM NaOH 0.2 mM Na_2_EDTA) for 1 hour in 95 °C, and equal volume of neutralizing solution (40 mM Tris-HCl) is added afterward. The genomic preparation is diluted 1:20 prior to HRM.

### Drift-Diffusion model

To better understand the origin of the observed behavioral phenotypes during motion integration, we use computational modeling. This approach provides us with a more detailed characterization of the behavior and allows us to indirectly infer how a mutation affects specific components of the sensory-motor transformation algorithm.

Previous work indicated that a simple Drift-Diffusion model with a decision threshold can explain many aspects of the responses to the dot motion stimulus (**Fig. 3e, Fig. S3a**,**b**, and ^28,29^). The model is based on the temporal integration of noisy (*σ*) motion evidence with certain coherence levels. The integrator is leaky and, therefore, its signal increases slowly with a time constant (τ) when the motion stimulus starts and decays with the same dynamics when it stops. In the model, fish swim spontaneously with two different probabilities. When integrated motion evidence is below the decision threshold (T), animals swim forward with a probability of *p*_*below*_. When it is above the threshold, they make a turn with a probability of *p*_*above*_. Notably, none of the underlying five model parameters can be measured directly through behavioral experiments, requiring us to resort to an indirect method, a multi-objective fitting strategy.

### Multi-objective fitting algorithm

To uncover latent changes within the motion integrating and decision-making circuits, we modified an evolutionary multi-objective optimization technique that can find the same global minimum and the same parameter set over repeated optimization runs^40,41^ (**Fig. S3c)**. This approach has been used in the past to solve highly non-linear models that require multiple behavioral features to be optimized simultaneously^42,43^. One starts with a population of randomly chosen parameter sets (800 individuals). Each parameter set gets evaluated, producing five behavioral features: 1) The probability to turn in the correct direction as a function of coherence (**Fig. 3b**). 2) The interbout interval as a function of coherence (**Fig. 3c**). 3) The probability to turn in the same direction for consecutive swims during 0% coherence (**Fig. 3d**). 4) The binned probability to turn in the correct direction as a function of time and coherence (**Fig. S2b**). 5) The turn angle probability distribution (**Fig. S2c**). For each of those features, the algorithm computes the squared distance to the same feature obtained in the experiments. Using the resulting five distance functions, the multi-objective algorithm then chooses which individuals are mutated and exchange parameter information using crossover to build the next generation.

We tested the validity of this approach in artificially created surrogate data sets (**Fig. S3d–h**), where we explicitly coded in certain hidden variables. The extent of their successful recovery allows for a quantitative evaluation of the multi-objective optimization technique. Using a set of parameters closely following our recently hand-tuned parameter set^28^, we find that the five extracted behavioral features generally capture what we find in the experiment (compare **Fig. S3d, Fig. 3b–d**, and **Fig. S2b**,**c**). After a few generations, the optimization algorithm was able to identify individuals that had near-zero error in one of the behavioral features. As the five distance functions have different scales, we normalized them using the 75 percentile of the distribution of error values. Finally, to assign a compromise error value to each individual, we computed a weighted sum of the five normalized distance functions. As we consider the interbout interval as the most important behavioral feature that our model should definitely capture, we set the weight for the interbout interval distance function to 3, for all other distance functions we used a weight of 1.

It took about 80 generations for this compromise error function to converge (**Fig. S3e**). Notably, our algorithm does not find individuals with exactly zero error. This is expected as our simulation is stochastic and, therefore, does not produce the exact same behavior in every stimulation run. We repeated the optimization algorithm 12 times for 4 models with different parameter sets (**Fig. S3f**). For each of the runs and models, we find that the algorithm successfully reduces the 5 error functions as well as the compromise error. At the end of each optimization run, we then picked the one individual with the smallest compromise error value and compared its parameter values to the parameters originally used to create the surrogate data set (**Fig. S3g**). We find that our algorithm can indeed reveal these values and that repeated optimization runs produce more or less the same results. Looking at one of those optimized models, we confirm that the behavioral features do indeed closely resemble the ones from the original dataset set (**Fig. S3h**). In summary, we conclude that our multi-objective evolutionary optimization is capable of extracting the hidden variables in our motion integration and decision-making model.

### Drift-diffusion model parameters extracted from motion integration assay

Knowing that our multi-objective fitting algorithm can reveal the latent variables in our drift-diffusion model, we next applied this strategy to real experimental behavioral data of larval fish with different genotypes. For all tested genotypes, the optimization algorithm produces behavioral dynamics closely mimicking the ones found in experiments (compare **Fig. 3b–d** and **Fig. 3f–h**, as well as **Fig. S2b**,**c** and **Fig. S2e**,**f**). We repeated the optimization algorithm 12 times for each dataset. We find solutions to have similarly small error values (**Fig. S4a**) and that the estimated model parameters are more or less identical after each run (**Fig. S4b**).

### Collective behavior model

We simulated groups of five fish freely swimming in a circular arena. The arena was modeled with a diameter of 1024 pixels. Each fish had a random starting position (x and y) and orientation in the arena and length of *fishsize* = 20 *pixels*. At every time point during the simulation, fish made swimming decisions based on two simple sensory-motor transformation rules: the tendency to avoid or approach clutter, and the tendency to align with global motion drift.

To compute clutter, we determined how much space another fish (*fish i*) occupies in the visual field of the focal fish (*fish j*) and added all values from the right hemisphere, and subtracted all values from the left hemisphere:

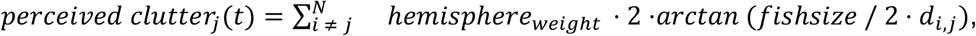

where 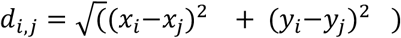 is the euclidean distance between animal pairs and where *hemisphere*_*weight*_ was either +1 or -1, depending on whether *fish i* and was in the right or left hemisphere relative to the orientation of *fish j*. N is the number of fish (= 5 fish in our simulations). Hence, the sign and amplitude of the resulting signal reflect the asymmetry perceived by the focal fish.

To compute the perceived motion by the focal fish (*fish j*), we used the perpendicular movement speed vectors of all other fish (*fish i*) relative to its own body orientation (*α*_*i,j*_, the angle between orientation vectors), linearly normalized by the euclidean distance (*d*_*i,j*_) between fish:

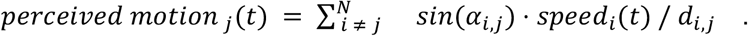

For example, when another fish approaches the focal fish at 90°, one gets a negative value, when it swims parallel, one gets zero, and when it swims away at an angle of 90°, one gets a positive value.

To obtain the total momentary perceived evidence for *fish j*, we simply added the values for clutter and motion:

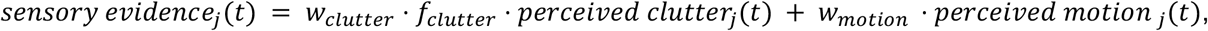

Where *w*_*clutter*_ is the weight of the clutter system, where *f*_*clutter*_ is the genotype- and age-dependent factor of the clutter system, and where *w*_*motion*_ is the weight of the motion system.

Following our drift-diffusion model (**Fig. 3e**), we integrated this value over time. Accordingly, when the integrated value was between the positive and negative thresholds, animals swam forward, when it was above the positive threshold, they turned to the right, and when it was below the negative threshold, they turned to the left. When swimming forward, animals covered 20 pixels per bout (about one body length) along their momentary axis of orientation. Moreover, animals stochastically changed their body orientation by a bit. Following the experimentally measured heading angle change distributions (**Fig. S2c**), we drew those angles from a Gaussian distribution centered at 0° with a standard deviation of 5°. Similarly, for initiating right or left turns, we drew angles from a Gaussian distribution centered at 22° or -22°, respectively, with a standard deviation of 25° (**Fig. S2c**). To capture the length and short gliding phase of forward swims and turns, we applied a low-pass filter with a time constant of 100 ms to these events, resulting in bout-like animal movement.

Notably, *w*_*clutter*_ and *w*_*motion*_ are the only free parameters of our model, all other parameters are directly extracted from experimental data. Even though tweaking those values would likely have resulted in overall improved model performance, we wanted to work with the most minimal model and, therefore, simply set both weights to 1. We further chose *f*_*clutter*_ for the 7 dpf *scn1lab*_*allele2*_^*+/+*^ wildtype to be -1, which reproduced the weak avoidance and alignment of 7 dpf *scn1lab*_*allele2*_^*+/+*^ wildtype larvae. The values for *f*_*clutter*_for the other genotypes and ages were then scaled according to the experimentally measured probability to turn towards clutter (**Fig. 2e**,**f** and **Fig. S1e**. For example, the clutter response strength for the 7 dpf *scn1lab*_*allele2*_^*+/-*^ was twice as strong as the one for the 7 dpf *scn1lab*_*allele2*_^*+/+*^. Hence, for this genotype, we scaled *f*_*clutter*_to -2. The clutter response of 21 dpf *scn1lab*_*allele2*_^*+/+*^ fish was positive and about three times as strong in amplitude as the one for 7 dpf *scn1lab*_*allele2*_^*+/-*^ larvae. Hence, we scaled this value to +3. This procedure led to the following clutter response factors for the 7 dpf animals: *scn1lab*_*allele2*_^*+/+*^: –1; *scn1lab*_*allele2*_^*>+/-*^: –2; *scn1lab*_*allele1*_^*+/+*^: –1; *scn1lab*_*allele1*_^*+/+*^: –1.5; *disc*^*+/+*^: –2; *disc*^*-/-*^: –1.5. For the 21 dpf animals, we obtained: *scn1lab*_*allele2*_^*+/+*^: +3; *scn1lab*_*allele2*_^*>+/-*^: +3; *scn1lab*_*allele1*_*+/+*: +3; *scn1lab*_*allele1*_*+/+*: +2; *disc*^*+/+*^: +1; *disc*^*-/-*^: +2.

For all other parameters of our model (time constant, **τ**; noise, ***σ***; decision threshold, **T**; swim probabilities below and above the threshold, ***p***_***below***_ and ***p***_***above***_), we chose exactly the results obtained from the multi-objective fitting procedure (**Fig. S4b**). For testing the clutter or motion systems in isolation (**Fig. 4c**,**d**), we simply set the weight of the respective other system to zero. Every time when an animal reached the circular border of the arena, we picked a new random orientation vector. We did not reflect or wrap trajectories across the border. We simulated the collective behavior model for 600 s with a timestep of dt = 0.01 using the forward Euler method. We used Python 3.8 and the real-time compiler numba. We stored the resulting trajectories in the exact same format as used for our experimental data, allowing us to use the same analysis scripts to extract values for aggregation and alignment (**Fig. 1**). The simulation source code is available online: (https://github.com/arminbahl/mutant_zebrafish_behavior).

## Supplementary Information

**Table 1:**
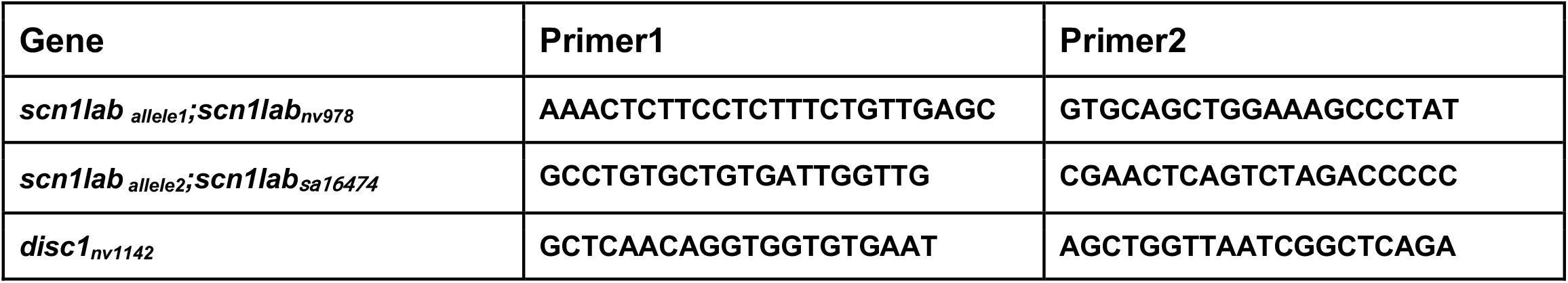
Genotyping Primers

**Figure S1.**
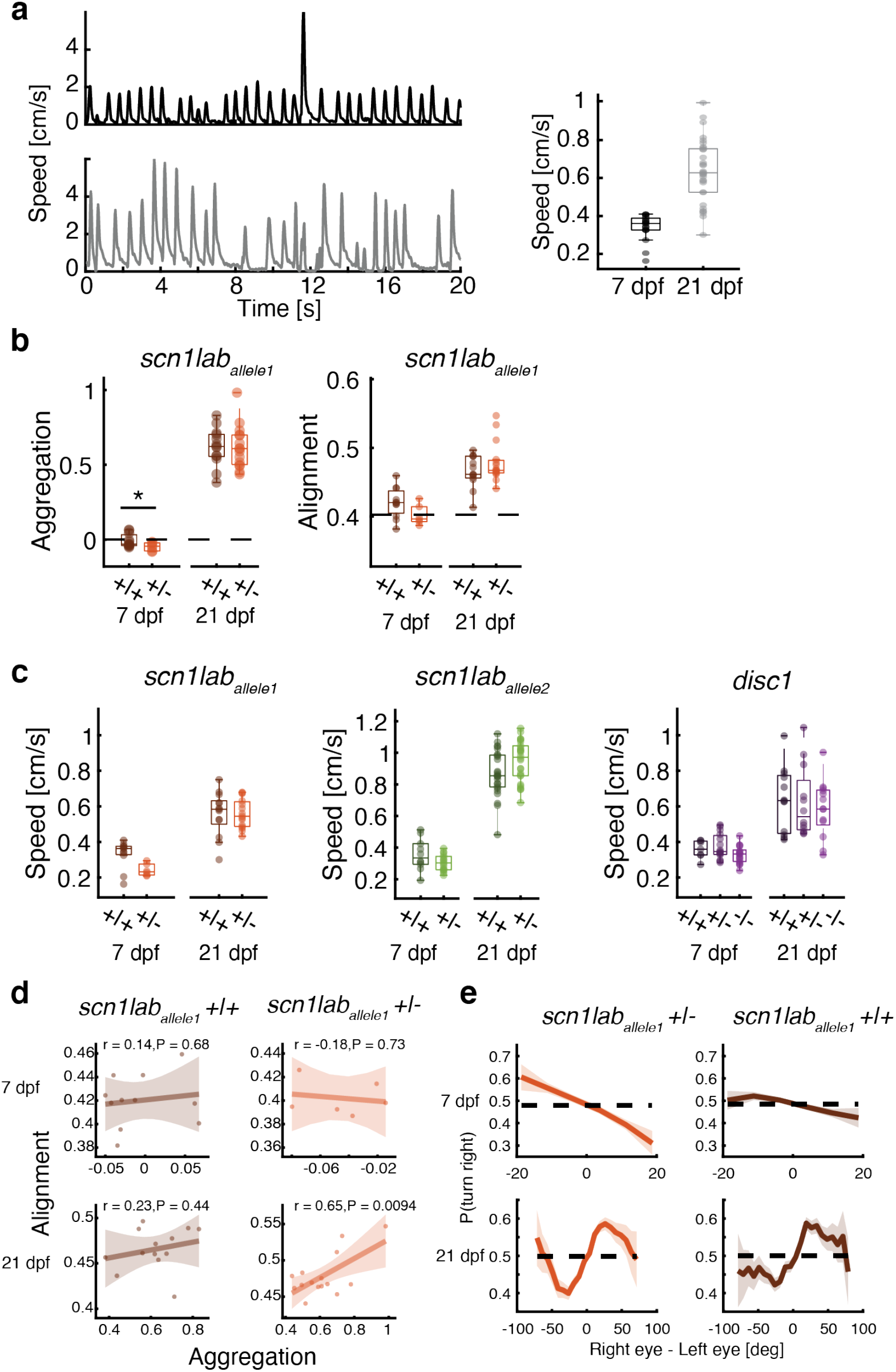
Single gene mutations do not affect speed during group assay. **(a)** Larval and juvenile zebrafish swim in bouts, with an overall increase in speed for older animals. (**b**) A mutation in the *scn1lab* gene (allele1) leads to more dispersed (less aggregated) groups, only in 7 dpf larvae (P=0.06, N = 15 groups; two-sided t-test) and has no effect on group alignment. (**c**) Compared to sibling controls, speed is not different in any of the tested mutants, but it is always larger in older animals. **(d)** Correlation of alignment and aggregation in *scn1lab*_*allele1*_ 7 (top row) and 21 dpf (bottom row) fish. No correlation is observed at 7 dpf (top), while 21 dpf Mutant *scn1lab*_*allele2*_^*+/-*^ fish (bottom right) show a stronger correlation, similar to the results seen in *scn1lab*_*allele1*_^*+/-*^ (**Fig. 1g**). **(e)** At 7 dpf s*cn1lab*_*allele1*_^+/-^ fish show a significant increase in the tendency to turn away from visual clutter, whereas at 21 dpf *scn1lab*_*allele2*_^+/-^ fish show a slightly reduced tendency to turn towards high visual clutter.

**Figure S2.**
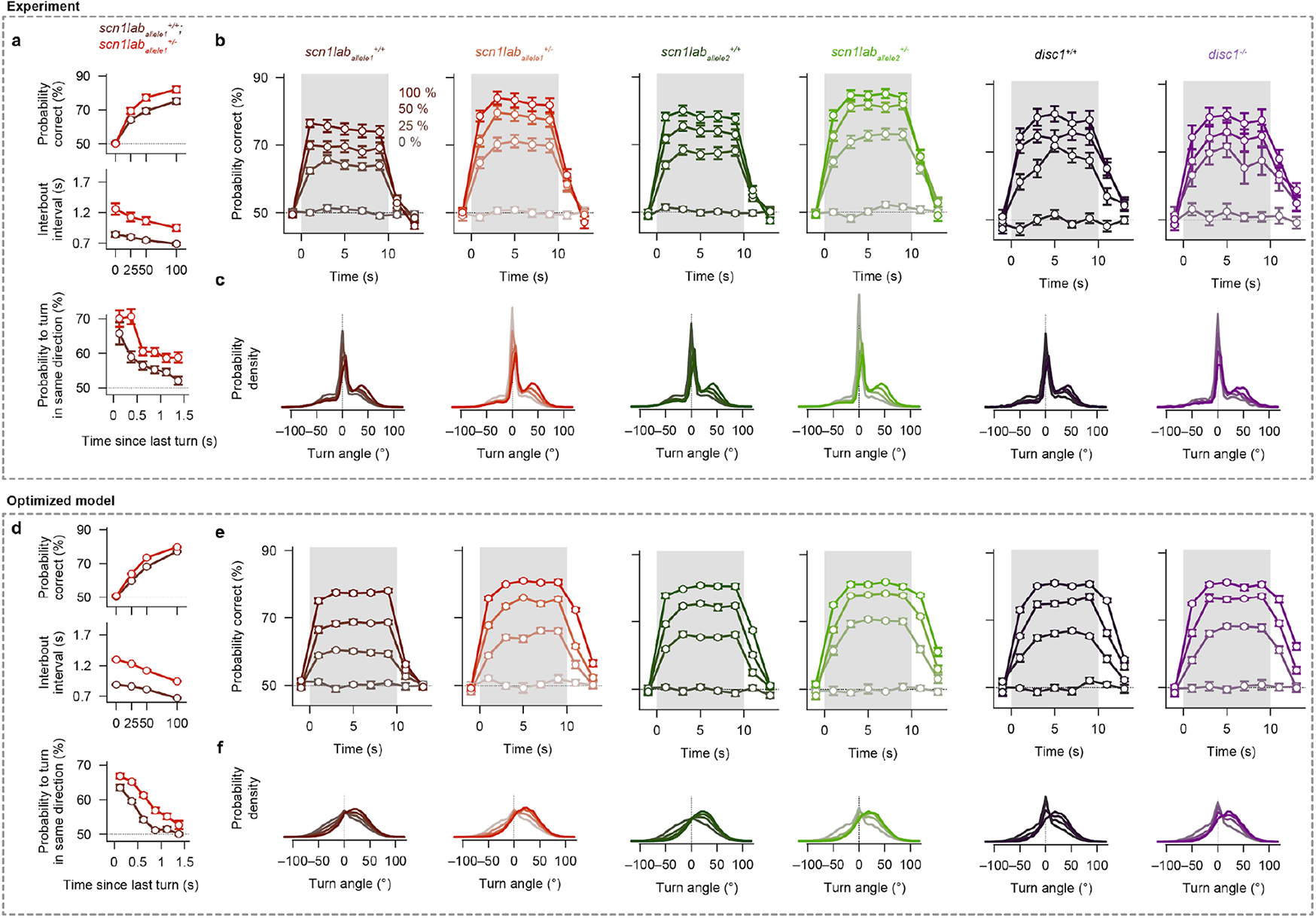
More detailed behavioral features and modeling results for the random dot motion integration assay for 7 dpf larvae. **(a)** The same analysis of experimental data as in **Fig. 2b–d** but for *scn1lab*_*allele1*_ mutant fish. The phenotypes of a mutation in this allele quantitatively match the ones for *scn1lab*_*allele2*._ (**b**) Probability correct over time for different coherence levels shows that both *scn1lab*^*+/-*^ mutants have an increased average probability correct and slower integration dynamics compared to *scn1lab*^*+/+*^ sibling controls. For *disc1*^*-/-*^ mutants, the response dynamics relative to *disc1*^*+/+*^ sibling controls does not change. (**c**) Turning angle probability density distribution shows that *scn1lab*^*+/-*^ mutant animals have a slightly higher tendency to make turns (30°-60°) than wild-type control siblings. For *disc1*^*-/-*^ mutants and their sibling controls, this effect is less pronounced. (**d–f**) Same analyses as in (**a–c**) but for the optimized models (same models as in **Fig. 2f–h**). In general, the model captures the phenotypes for *scn1lab*_*allele2*_ *and* the overall dynamics of the probability correct as a function of time. It also qualitatively reproduces the turning angle probability density distributions. Color saturation indicates coherence level (from less saturated to more saturated: 0%, 25%, 50%, and 100%). Darker colors indicate sibling controls. Red and green lines are *scn1lab*^*+/-*^ fish, respectively. Violett lines indicate *disc1*^*-/-*^ fish. N = 34, 27, 44, 36, 21, and 16 fish for genotypes *scn1lab*_*allele1*_^*+/+*^, *scn1lab*_*allele1*_^*+/-*^, *scn1lab*_*allele2*_^*+/+*^, *scn1lab*_*allele2*_^*+/-*^, *disc1*^*+/+*^, *and disc1*^*-/-*^, respectively, in **a–c**. N = 12 models (different optimization repeats) for each genotype in (**d–f**). All error bars are ±sem. Same fish and models as in **Fig. 2b–d,f–h**.

**Figure S3.**
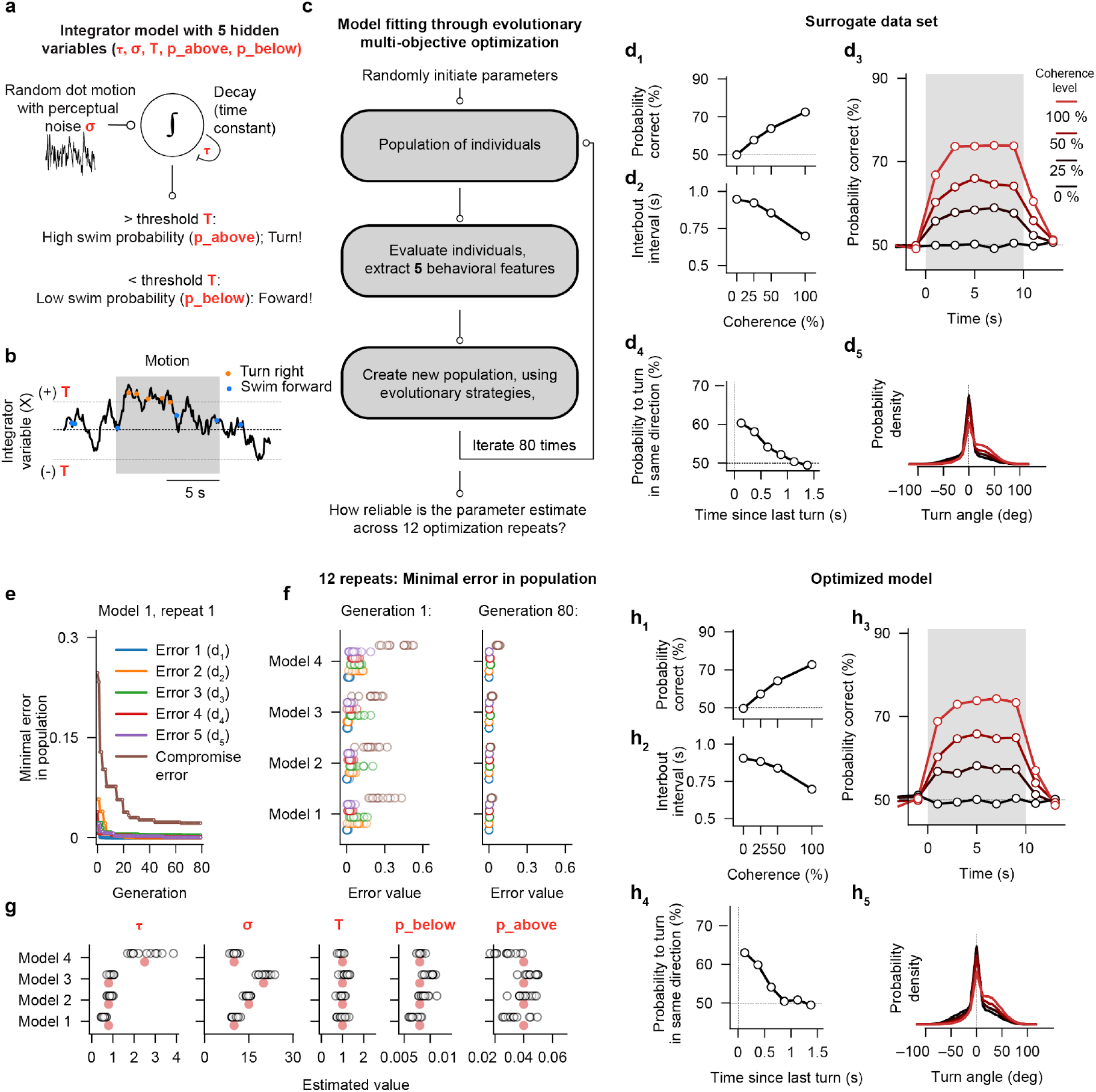
Multi-objective optimization strategy on surrogate data. (**a**,**b**) Schematic of the integrator and decision-making model. Motion evidence with perceptual noise (σ) is integrated by a leaky integrator with a time constant (τ). When the integrated value is below the decision threshold (T), the model creates forward swims with a probability of p_below_ (blue dots). Otherwise, it makes turns (orange dots) with a probability of p_above_. (**c**) Flow diagram of the evolutionary multi-objective optimization strategy. The algorithm starts with a population of randomly chosen individuals (parameter sets) and uses evolutionary principles to iteratively propagate models across generations. Models are chosen based on 5 multi-objective behavioral features, without needing to weigh them. We determine the general reliability and quality of the fitting algorithm by trying to uncover a few hidden model parameter combinations used to create artificial surrogate datasets. (**d**_**1**_**–d**_**5**_) The five behavioral features used as error functions during the multi-objective optimization (same analysis as in **Fig. 2b–d** and **Fig. S2b**,**c**). These example traces were created using the manually chosen parameter set given by the red dots for model 1 in panel **f**. (**e**) Evolution of error over generations for the 5 error functions. After a few generations, these error values converge to nearly zero. The compromise error (a range-corrected weighted sum of all 5 error functions, see methods) requires more generations to converge. (**f**) Minimal error for the 5 error functions and the compromise error for the first and last generation for 12 optimization repeats and 4 example surrogate data sets. In the last generation, all 12 optimization repeats lead to almost identical error values. (**g**) Estimated parameters for the 4 example models. Optimization repeats are indicated with open black circles. The target parameter used to create the surrogate dataset is indicated with the red circle. The optimization algorithm can reliably reveal the hidden variables. (**h**_**1**_**–h**_**5**_) Simulation results of the optimized model following parameter optimization using the surrogate dataset of model 1 as a target (**d**_**1**_**–d**_**5**_). The objectives between the two simulations precisely match.

**Figure S4.**
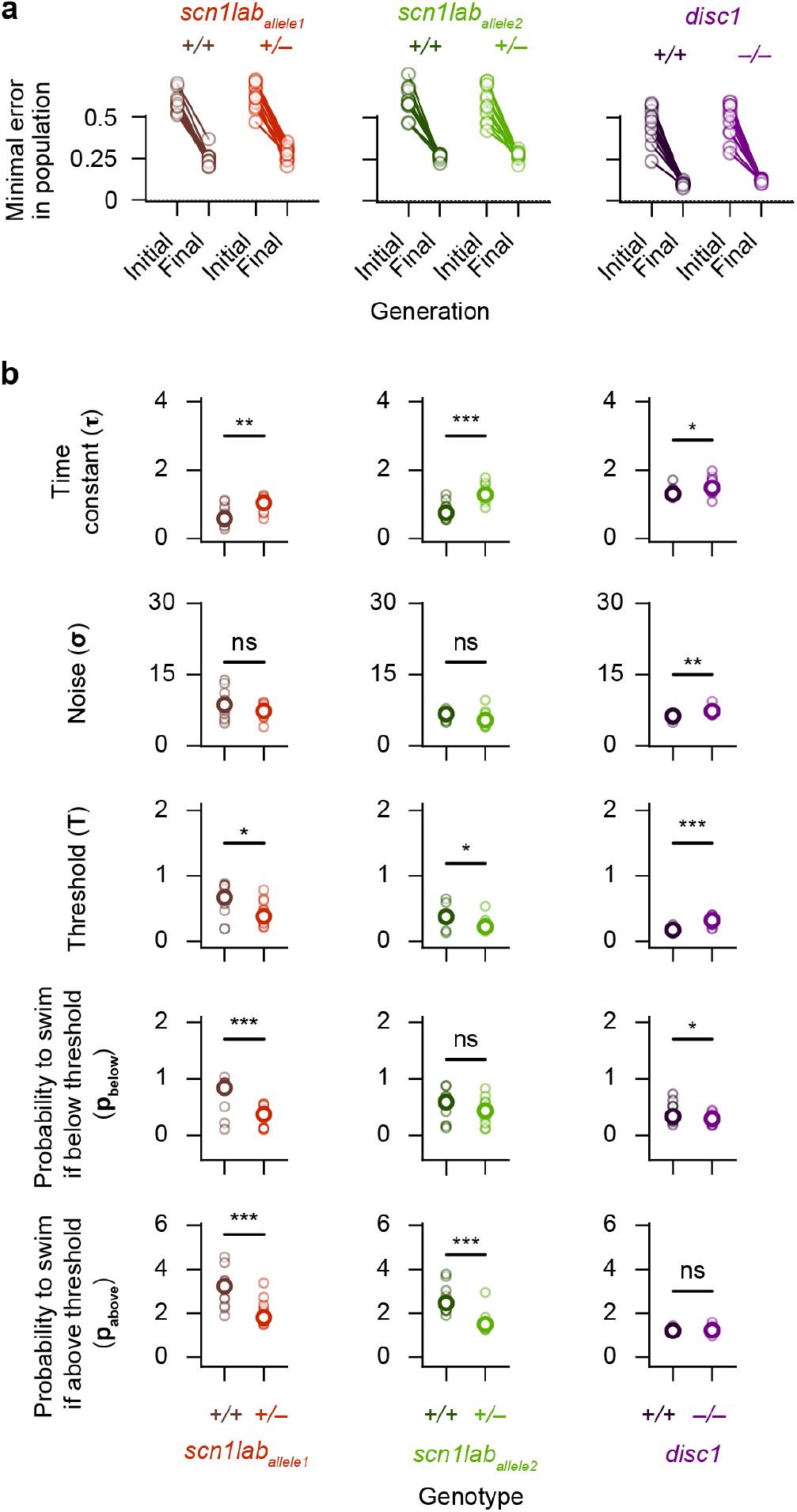
Mutant larval zebrafish have specific algorithmic alterations in their ability to integrate motion signals. **(a)** Value of the compromise error function of the best individual in the initial generation and after 80 generations. 12 optimization runs and 6 target genotypes (3 mutants with respective sibling controls). **(b)** Estimated model parameter values of the best individual for each genotype and optimization run *p < 0.05, **p < 0.01; ***p < 0.001; calculated by comparing the actual difference between the parameter median of each genotype to the distribution of differences for 10000 bootstrap repeats.) N = 12 model optimization repeats. Small open circles are individual repeats, bigger open circles are median parameter values.

**Video S1: Freely swimming larval zebrafish in the random-dot-motion integration assay**. The visual scene, projected from below, consists of a cloud of randomly flickering small dots. A fraction of those dots moves coherently either to the left (left panel) or to the right (right panel), relative to the body orientation of the animal. In this video, we illustrate the 100 % coherence stimulus. For lower coherence values, fewer dots move. The motion direction is constantly updated in closed-loop, creating uni-directional motion flow from the perspective of the fish. Fish integrate this information over time and use these cues to make swimming decisions.

**Video S2: Model simulation results for 7 dpf wild-type larvae**. We simulated a group of five fish swimming in a circular arena (**Fig. 4b**, left panel). These young fish slightly swim away from clutter (**Fig. 1e**, middle panel, and **Fig. 4a**), and integrate all motion cues in their environment (**Fig. 3a**,**e** and **Fig.4a**). In this configuration, the virtual group is slightly over-dispersed (less aggregated than expected by chance, Fig. 4c, left panel) and animals minimally align with each other (**Fig. 4d**, left panel).

**Video S3: Model simulation results for 21 dpf wild-type fish**. We simulated a group of five 21 dpf fish swimming in a circular arena (**Fig. 4b**, right panel). Here, clutter is attractive (**Fig. 1e**, rightmost panel, and **Fig. 4a**). The model parameters for motion integration and decision-making are the same as for 7 dpf fish. In this configuration, we see an emergence of group behaviors: The virtual group shoals (animals are more aggregated than expected by chance, **Fig. 4c**, right panel) and fish robustly align with each other (**Fig. 4d**, right panel).

